# Loud acoustic stimulation reveals an online reticulospinal contribution to long-latency reflexes in humans

**DOI:** 10.64898/2026.08.10.743894

**Authors:** Hirotaka Sugino, Daichi Nozaki, Junichi Ushiyama

**Affiliations:** Graduate School of Media and Governance, Keio University, Kanagawa, Japan; Graduate School of Education, The University of Tokyo, Tokyo, Japan; Faculty of Environment and Information Studies, Keio University, Kanagawa, Japan; Department of Rehabilitation Medicine, Keio University School of Medicine, Tokyo, Japan

**Keywords:** reticulospinal tract, long-latency reflex, loud acoustic stimulation, corticospinal tract, human motor control

## Abstract

The long-latency reflex (LLR), the fastest feedback response that recruits supraspinal pathways, is an important model for understanding how descending motor pathways shape rapid corrective responses in humans. While the corticospinal tract’s contribution to the LLR has been well established, that of the reticulospinal tract, the other major descending motor pathway, remains purely speculative. To address this online contribution to the generation of the LLR, we used loud acoustic stimulation (LAS), which can strongly engage brainstem circuits including the pontomedullary reticular formation. By delivering LAS at nine timings (0–80 ms in 10-ms steps) relative to perturbation onset, we tested whether LAS selectively facilitates the LLR but not the short-latency reflex (SLR), and whether the facilitated epoch shifts systematically with LAS timing. In twelve healthy participants, elbow extension perturbations were applied to evoke stretch reflexes in the biceps brachii muscle. LAS produced significant supralinear facilitation in the LLR but not in the SLR. Moreover, at LAS timings of 50 ms or more after perturbation onset, LLR facilitation shifted progressively later with LAS, remaining at an approximately fixed delay of 30 ms after LAS onset. This fixed delay indicates that LAS-evoked descending input from the same origin facilitates the ongoing LLR. Together with the lack of significant SLR facilitation, this temporal pattern supports an online reticulospinal contribution to the human LLR, alongside the established corticospinal contribution. This approach provides a new, non-invasive means to investigate the physiological role of the reticulospinal tract in human motor control.

**Key Points:**

- The long-latency reflex is a rapid muscle response to sudden stretch. Unlike faster spinal reflexes, it is shaped by commands descending from the brain and adjusts to the task.
- Though the corticospinal tract is known to shape this reflex, whether the reticulospinal tract also contributes to the reflex has not been tested in humans.
- We stretched the arm and, at various delays, played a loud sound that engages the brainstem origin of the reticulospinal tract. The sound significantly enhanced the long-latency reflex, whereas no significant enhancement was detected in the faster spinal reflex.
- When the sound came 50 milliseconds or more after the stretch, the enhancement followed the sound at a stable delay, indicating that sound-evoked descending signals interacted with the ongoing reflex response.
- These findings support a real-time contribution of the reticulospinal tract to the human long-latency reflex and provide a non-invasive way to study this pathway.

## Introduction

The long-latency reflex (LLR) provides a critical model for understanding how descending motor pathways shape rapid corrective responses in humans. Muscle stretch evokes two principal reflex components: the short-latency reflex (SLR, 20–50 ms) generated by spinal circuits and the LLR (50–100 ms) that recruits supraspinal pathways and is highly sensitive to task demands (Kurtzer, 2015). Among descending motor pathways, the contribution of the corticospinal tract to the LLR has been firmly established in humans (Kurtzer, 2015). For example, transcranial magnetic stimulation (TMS) studies have shown that corticospinal drive facilitates electromyogram (EMG) activity during LLR, but not during the SLR (Day et al., 1991; Palmer and Ashby, 1992). This evidence supports the view that the LLR is shaped, at least in part, by corticospinal processing, whereas the SLR reflects spinal processing alone.

The reticulospinal tract, the other major descending motor pathway in primates, is also assumed to contribute to the LLR (Shemmell et al., 2010; Kurtzer, 2015). This assumption is supported by converging but largely indirect lines of evidence. Anatomically, proprioceptive input from the limbs can reach the reticular formation (Leiras et al., 2010), and reticulospinal fibres are known to project prominently to the spinal intermediate zone (Lawrence and Kuypers, 1968a, b). Physiologically, recordings in animals have shown that pontomedullary reticular neurons discharge rapidly in response to limb perturbations (Stapley and Drew, 2009) and receive converging limb somatosensory and vestibular inputs (Miller et al., 2017). In humans, functional neuroimaging has revealed brainstem activation during stretch reflexes (Zonnino et al., 2021). However, these approaches could not provide temporally precise assessments of whether and how activation of the reticulospinal tract contributes to the LLR on a millisecond timescale. A major limitation in elucidating this contribution is the lack of a non-invasive approach that can probe reticulospinal drive within the brief time window of the stretch reflex. Thus, despite being a major descending motor pathway, the real-time contribution of the reticulospinal tract to the human LLR remains unclear, leaving its role in rapid corrective responses an open question.

Loud acoustic stimulation (LAS) may provide a means of overcoming this limitation. LAS strongly engages startle-related brainstem circuits, including the pontomedullary reticular formation, and has been used to examine reticulospinal contributions to movement through the generalised startle response (Brown et al., 1991; Yeomans et al., 2002; Zheng and Schmid, 2023) and the StartReact phenomenon (Valls-Solé et al., 1999; Nonnekes et al., 2014; Tapia et al., 2022). Extending this approach to stretch reflexes, Foysal et al. (2016) demonstrated that repeatedly pairing LAS with muscle stimulation can induce offline plasticity in the LLR, suggesting that reticulospinal drive can influence the stretch reflex circuit. However, whether this drive shapes LLR output online has not been directly tested.

Here, we developed a non-invasive approach that combines mechanical perturbation with LAS to investigate the online contribution of the reticulospinal tract to the generation of the LLR in humans. To this end, we adapted the supralinear summation logic originally developed for TMS (Day et al., 1991; Palmer and Ashby, 1992; Lewis et al., 2004, 2006; Pruszynski et al., 2011; Perenboom et al., 2015). LAS was applied at nine timings (0–80 ms in 10-ms steps) relative to perturbation onset and tested whether LAS-evoked reticulospinal drive produces supralinear facilitation specifically within the LLR window, leaving the SLR unaffected. Additionally, we tested whether the facilitated epoch shifts together with changes of the timing of LAS onset. If that is the case, we could consider the observed facilitation of the LLR to be temporally linked to LAS-evoked descending input. The results provide evidence supporting the online contribution of the reticulospinal tract to the human LLR.

## Materials and Methods

### Ethical approval

This study was approved by the Research Ethics Committee of Saiseikai Higashikanagawa Rehabilitation Hospital (Approval Number 20-12) and the Research Ethics Committee of the Shonan Fujisawa Campus (SFC), Keio University (Approval Number 506). Written informed consent was obtained from all participants prior to the experiment. All research procedures were performed in accordance with the guidelines established in the Declaration of Helsinki, except that the study was not preregistered in a database.

### Participants

Twelve healthy individuals (age, 23.0 ± 2.5 years [mean ± SD]; 6 males and 6 females) participated in this study. All participants were right-handed and had no history of neurological, musculoskeletal and hearing disorders.

### Apparatus

This study used a robotic exoskeleton (KINARM Exoskeleton; KINARM, Kingston, Canada) that permits participants to flex and extend the shoulder and elbow joints in the horizontal plane and can impose mechanical loads independently at the shoulder and/or elbow. Visual targets and a cursor representing hand position were projected onto a horizontal mirrored display positioned above the arm, which also occluded direct vision of the limb. Joint angles were sampled at 1 kHz.

### EMG recording

Surface EMG signals were recorded from the right biceps brachii (BB), the lateral head of the right triceps brachii (TB), and the right sternocleidomastoid (SCM). SCM activity was recorded as a marker of the acoustic startle reflex. Indeed, SCM has been commonly used as an indicator of the overt startle response to auditory stimuli in humans (Brown et al., 1991; Valls-Solé et al., 1999). For each muscle, two disposable electrodes (Kendall H124SG; Cardinal Health, Dublin, OH, USA) were attached directly to a wireless Pico EMG sensor (Wave Plus system; COMETA, Milan, Italy) and positioned over the muscle belly. The inter-electrode distance was fixed by the sensor geometry at 25 mm. Signals were amplified (gain ×1000) and band-pass filtered (10–500 Hz). The analogue output was fed to the KINARM system for synchronised acquisition. All EMG signals were sampled at 1 kHz.

### Loud acoustic stimulation

The LAS consisted of a 500-Hz tone burst delivered at 110 dB SPL for 50 ms from a loudspeaker (YAMAHA DBR15, Yamaha Corporation, Hamamatsu, Japan) placed 30 cm posterior to the participant’s head. This stimulation profile is known to activate the reticular formation (Yeomans et al., 2002; Mooney et al., 2023). Before data collection, participants were exposed to five auditory presentations to familiarise them with the LAS and were asked to report any discomfort. All participants described the stimulus as tolerable and non-painful. Startle-like movements were commonly visible during the initial presentations but diminished over subsequent presentations, consistent with rapid habituation (Brown et al., 1991).

### Procedures

To investigate whether the reticulospinal tract is involved in the stretch reflex, we conducted an experiment combining mechanical perturbation with LAS by adapting the approach used in studies of corticospinal involvement in stretch reflexes using TMS (Day et al., 1991; Palmer and Ashby, 1992; Lewis et al., 2004, 2006; Pruszynski et al., 2011; Perenboom et al., 2015). Participants sat in the KINARM exoskeleton and controlled a cursor on the screen with their right hand by gripping and manipulating the handle. After task initiation, participants aligned the cursor (0.8 cm diameter) with a target circle (1 cm diameter) at a posture of 40° elbow flexion and 70° shoulder flexion, against a 1.5 Nm elbow flexion background load. After a randomised interval between 3 and 7 s, one of eleven trial types was presented in a random order (Fig. 1): the LAS-only condition, the perturbation-only condition, or the perturbation-and-LAS conditions with nine different timings between LAS and mechanical perturbation (see below).

**Fig 1.**
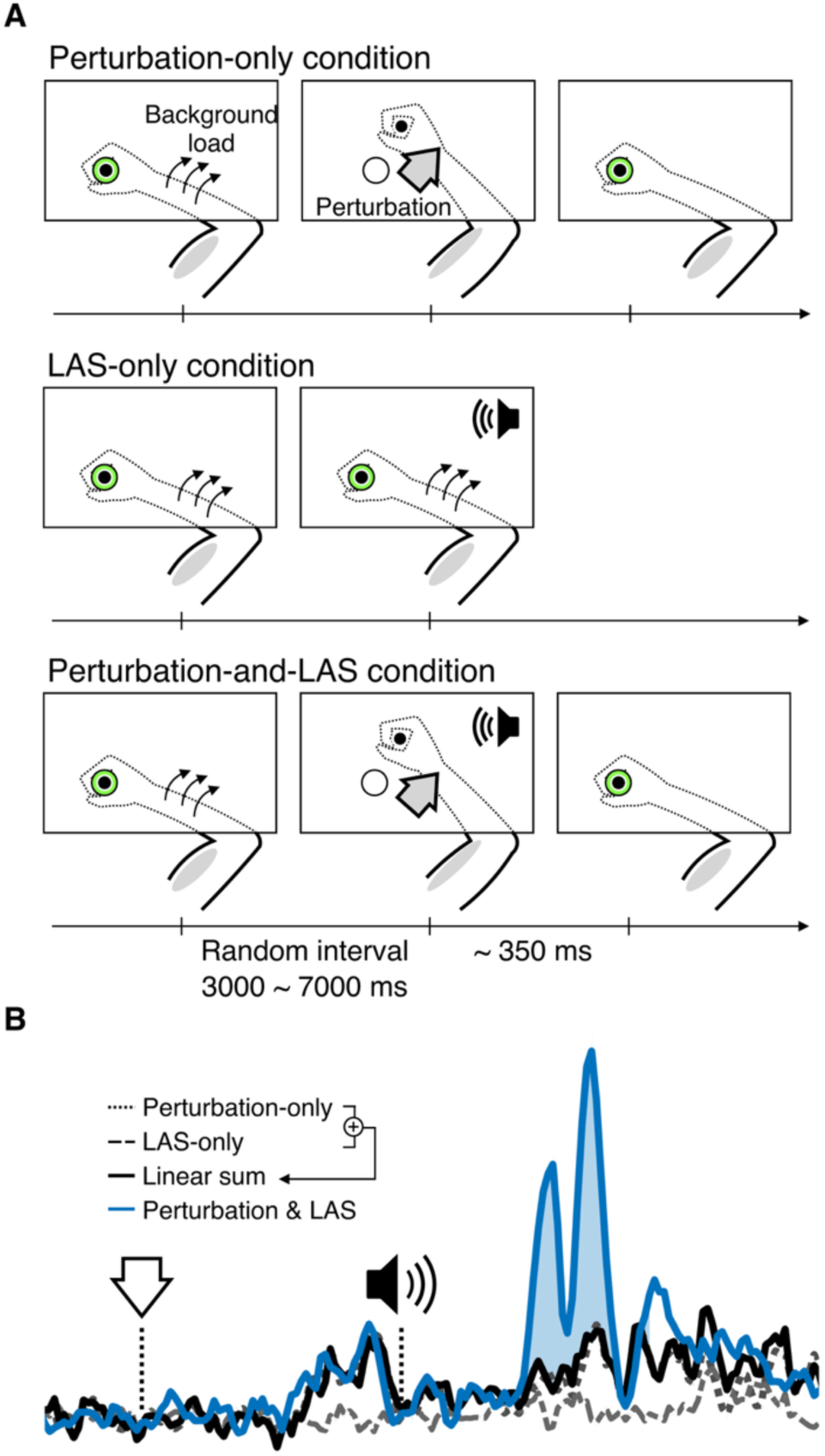
Experimental paradigm and supralinear summation logic. **(A)** Participants held a handle connected to KINARM exoskeleton and maintained their start posture (40° elbow flexion, 70° shoulder flexion) against a 1.5 Nm elbow flexion background load, aligning a cursor (0.8 cm diameter) with a target (1 cm diameter). After a randomised 3–7 s interval, one of eleven trial types was presented: LAS-only, in which a 110 dB SPL, 50 ms, 500 Hz tone burst was delivered alone; perturbation-only, in which a 1.5 Nm elbow extension perturbation was delivered for 1 s; or one of nine perturbation-and-LAS conditions, in which the LAS was delivered at 0, 10, 20, 30, 40, 50, 60, 70 or 80 ms after perturbation onset. Each participant completed 20 trials per condition. **(B)** Schematic of the supralinear summation logic applied to a representative perturbation-and-LAS condition. The downward arrow indicates perturbation onset, and the speaker icon indicates LAS onset. The linear sum (black solid) is computed as the perturbation-only EMG (dotted) plus the time-shifted LAS-only EMG (dashed). Supralinear facilitation is inferred when the perturbation-and-LAS EMG (blue solid) exceeds the linear sum.

In the LAS-only condition, the LAS was delivered alone and the trial terminated 2 s after the LAS with removal of the background load. In the perturbation-only condition, a mechanical perturbation (1.5 Nm elbow extension, 1-s duration) displaced the cursor from the target, and then participants were instructed to return the cursor to the target as quickly as possible. After holding the target for 1 s, the background load was removed and the trial ended. Visual feedback (“Good” for return times ≤ 350 ms after perturbation onset, “Slow” otherwise) was displayed at the end of each perturbation trial. In the perturbation-and-LAS conditions, the LAS was delivered at one of nine timings relative to perturbation onset: 0, 10, 20, 30, 40, 50, 60, 70 or 80 ms. Similar to the perturbation-only condition, participants were instructed to return the cursor to the target as quickly as possible.

Participants first completed 10 practice trials in which trial types were randomly selected from the same eleven conditions used in the main experiment. In the main experiment, the 22 trials in each set comprised two repetitions of each of the eleven conditions and were presented in a computer-generated random order. Across 10 sets, this yielded 20 trials per condition per participant. Participants could rest freely between sets and took a mandatory break of at least 5 minutes after set 5 to minimise fatigue.

### Analysis

Data analysis was performed using custom scripts in MATLAB (MathWorks, Natick, MA, USA). EMG signals were notch-filtered at 49–51 Hz to remove line noise, band-pass filtered (20–450 Hz, second-order Butterworth, zero-phase), and full-wave rectified. For each trial, EMG signals were aligned to perturbation onset except LAS-only trials, which were aligned to LAS onset. Then they were normalised to the mean EMG in a 100-ms baseline window preceding perturbation onset, or LAS onset for LAS-only trials. Trials containing movement artefacts were rejected automatically by an amplitude criterion applied separately to each participant and each muscle: within a window spanning from the start of the baseline to the end of the LLR window (100 ms after perturbation onset), the peak rectified EMG was computed per trial, and trials whose peak exceeded five times the within-participant median peak were excluded. Artefact rejection was performed independently for BB, TB, and SCM, so each muscle retained its own valid trial set. Consequently, this criterion excluded one trial from the BB analysis and one different trial from the TB analysis. No trials were excluded from the SCM analysis. All twelve participants were included in all analyses. Normalised EMG were then averaged across trials within each condition.

We first performed three control analyses. For these control analyses, background EMG and SCM startle amplitudes were computed from the rectified and filtered EMG signals before baseline normalisation. To confirm that background muscle activity was similar across conditions, mean EMG in the 100-ms window preceding perturbation onset was compared across all eleven conditions, separately for BB and TB. To assess whether the initial perturbation kinematics differed among the ten conditions that included a mechanical perturbation (i.e., the perturbation-only condition and the nine perturbation-and-LAS conditions), we quantified elbow displacement 50 ms after perturbation onset and the maximum absolute elbow angular velocity from 0 to 50 ms (Vmax) from each participant’s condition-averaged trajectory. This early window was selected to characterise the imposed stretch before substantial mechanical effects of the reflex response could alter the trajectory (Tsuji and Rothwell, 2002). To assess whether the startle response persisted across the experiment, peak EMG amplitude of the SCM was compared between a 100-ms window 20– 120 ms after LAS onset and a 100-ms window before LAS in the LAS-only condition (pre-LAS baseline window). The choice of SCM and the 20–120 ms post-LAS evaluation window was informed by previous studies of human acoustic startle responses (Brown et al., 1991; Valls-Solé et al., 1999).

The main analysis for the facilitation of the stretch reflex was performed on BB, the muscle stretched by the perturbation. To quantify supralinear facilitation, we compared the observed BB EMG in each of nine perturbation-and-LAS conditions with the linear sum, computed as the perturbation-only EMG plus the LAS-only EMG time-shifted by the corresponding LAS timing. Because EMG was normalised to baseline, the common baseline was subtracted once to avoid counting it twice. This linear summation approach follows previous studies that probed corticospinal involvement in the LLR by combining TMS with the stretch reflex (Day et al., 1991; Palmer and Ashby, 1992; Lewis et al., 2004, 2006; Pruszynski et al., 2011; Perenboom et al., 2015).

For each participant and LAS timing, the linear sum was subtracted from the trial-averaged BB EMG waveform of the corresponding perturbation-and-LAS condition (observed − linear sum), yielding a facilitation waveform. Each facilitation waveform was smoothed with a 5-ms moving average along the time axis to attenuate high-frequency fluctuations in the rectified EMG and thereby obtain a stable estimate of the facilitation time course. This smoothed waveform was used both for the statistical analysis of significant facilitation and for the computation of the facilitation latency (below).

In addition, for each participant and LAS timing, we determined the onset of the facilitation using a threshold-crossing criterion adapted from Hodges and Bui (1996). The threshold was the mean of the facilitation waveform plus twice its standard deviation, both measured over the 100-ms window preceding perturbation onset. Searching forward from LAS onset to 129 ms after perturbation onset, the facilitation onset was taken as the first time point at which the smoothed facilitation waveform crossed this threshold and exceeded it for at least 5 consecutive ms. The facilitation latency was then defined as the interval from the timing of LAS onset to that of the facilitation onset.

### Statistical analysis

All statistical analyses were performed in MATLAB. Background EMG across all eleven conditions was compared using a one-way repeated-measures ANOVA, separately for BB and TB. When Mauchly’s test indicated violation of sphericity (*p* < 0.05), the Greenhouse– Geisser correction was applied to the degrees of freedom. On the other hand, when normality was violated (Shapiro–Wilk *p* < 0.05), the Friedman test was used in place of ANOVA. For the kinematic control analysis, elbow displacement at 50 ms and Vmax were each compared among the ten conditions that included a mechanical perturbation using a one-way repeated-measures ANOVA. The Greenhouse–Geisser correction was applied when Mauchly’s test indicated a violation of sphericity. The two omnibus *p* values were corrected using the Holm method. Comparisons between individual perturbation-and-LAS conditions and the perturbation-only condition were planned only when the corresponding Holm-corrected omnibus test was significant. For the SCM startle verification, the peak amplitude in the 20– 120 ms window after LAS onset was compared with that in the pre-LAS baseline window using a paired *t* test (or Wilcoxon signed-rank test when Shapiro–Wilk normality was violated).

The main facilitation analysis tested where, across post-perturbation time and LAS timing, the facilitation exceeded zero. For each participant, the smoothed facilitation waveforms were assembled into a two-dimensional map spanning post-perturbation time (0–129 ms) and the nine LAS timings (0–80 ms). Facilitation was tested against zero at every point of this map with a one-sample *t* test. Because the map contains more than a thousand points, significance was determined with a two-dimensional cluster-based permutation test rather than point by point. We followed the procedure of Maris and Oostenveld (2007), implemented with the permutest MATLAB function, which is provided with the analysis code. Adjacent points exceeding the threshold corresponding to *p* < 0.05 (one-sided) were grouped into clusters, and each cluster was summarised by the sum of its t statistics (T-sum). The null distribution of the T-sum was derived from the complete set of 4096 sign-flipping permutations (2^12^ for the twelve participants; Nichols and Holmes, 2001), and clusters yielding a permutation *p* < 0.05 were deemed significant. As this procedure controls the family-wise error rate across the entire map, no additional correction was applied. Because the tested map encompassed both the SLR and the LLR, facilitation could be evaluated across both predefined reflex windows.

The facilitation latency was analysed with a linear mixed model, with LAS timing as a categorical fixed effect (nine levels, reference 0 ms) and a random participant intercept, fitted by restricted maximum likelihood (in Wilkinson notation, latency ∼ LAS timing + (1 | participant)). This model used all available observations despite the data being unbalanced, because a facilitation onset was not detected for every participant at every LAS timing. The omnibus effect of LAS timing was tested with an *F* test using Satterthwaite-approximated degrees of freedom (Satterthwaite, 1946). From the same model, we obtained the estimated marginal mean latency and its 95% confidence interval (EMM) at each LAS timing, together with all 36 pairwise contrasts between timings, which were corrected for multiple comparisons using the Benjamini–Hochberg false discovery rate (Benjamini and Hochberg, 1995).

## Results

Figure 2 illustrates the group-averaged data for elbow joint displacement (A), and EMG signals from the BB (B) muscles in the perturbation-only condition. The elbow extension perturbation displaced the elbow joint by approximately 4°, reaching peak displacement at approximately 200 ms after perturbation onset and returning to near-baseline by 500 ms. As a result of the elbow extension perturbation, the BB EMG exhibited a clear SLR (20–50 ms) followed by a distinct LLR (50–100 ms), confirming that the perturbation reliably evoked a stretch reflex. In addition, Figure 2C illustrates the group-averaged SCM EMG in the LAS-only condition. Qualitatively, the SCM EMG showed no characteristic EMG activity following LAS onset.

**Fig 2.**
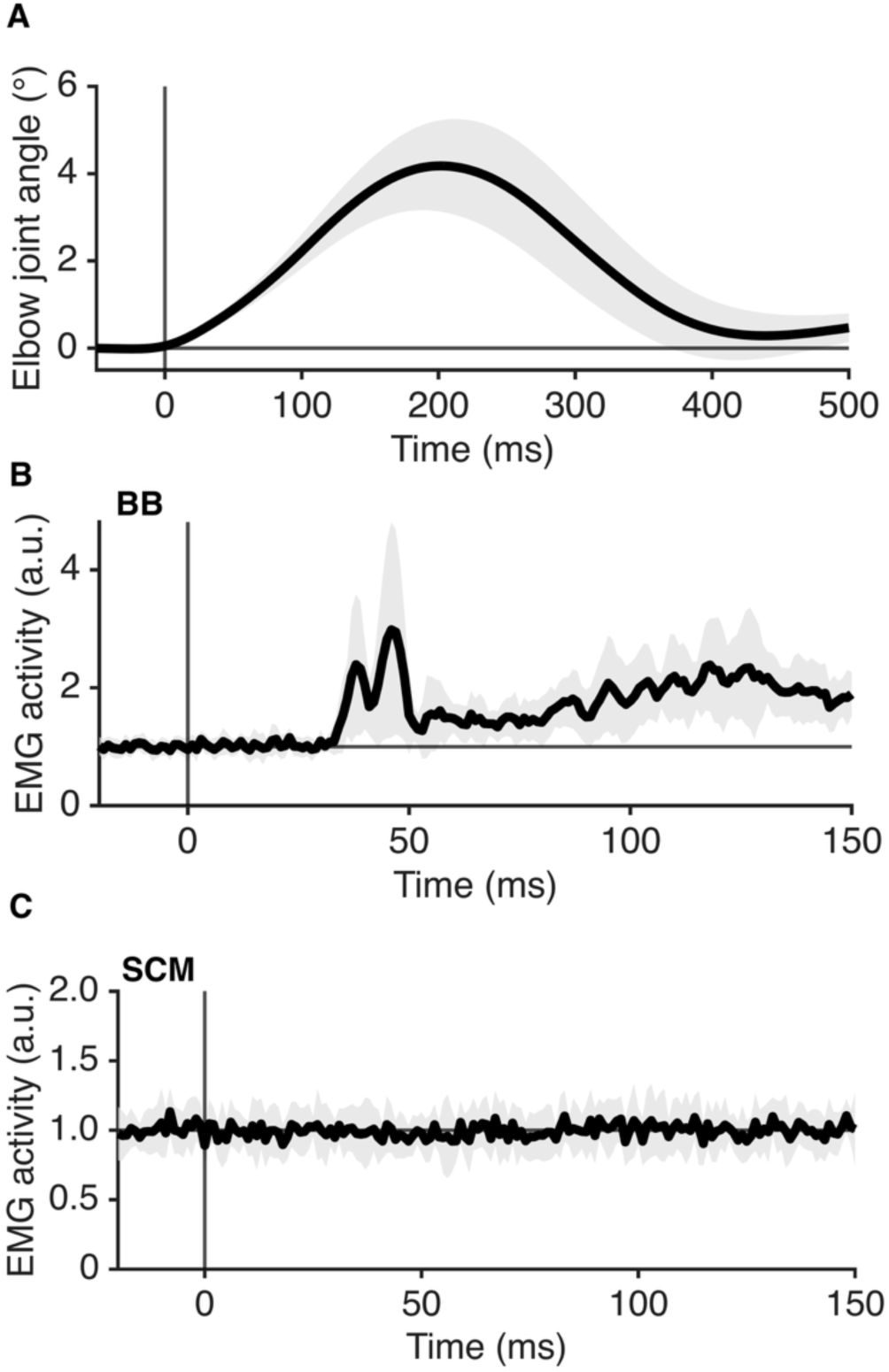
Perturbation-evoked elbow displacement, biceps EMG, and the sternocleidomastoid EMG response during the LAS-only condition. **(A)** Group-averaged elbow joint angular displacement in the perturbation-only condition. **(B)** Group-averaged biceps brachii (BB) EMG in the perturbation-only condition, including the short-latency reflex (SLR, 20–50 ms) and the long-latency reflex (LLR, 50–100 ms). **(C)** Group-averaged sternocleidomastoid (SCM) EMG in the loud acoustic stimulation (LAS)-only condition. The vertical line indicates perturbation onset (A–B) or LAS onset (C). Black line, mean; grey shading, ± SD. In (B) and (C), EMG is normalised to the mean baseline in the 100-ms window preceding perturbation onset (B) or LAS onset (C) and is expressed in these baseline-normalised units (a.u.); the horizontal line at 1 marks baseline activity.

To assess whether pre-perturbation background muscle activity varied across trial types, the mean background EMG measured from the 100-ms window before perturbation onset did not differ significantly across the eleven conditions for either the BB (F(10, 110) = 1.403, p = 0.188, ηp2 = 0.113) or the TB (F(10, 110) = 1.785, p = 0.0716, ηp2 = 0.140). These results provided no evidence of a condition-dependent change in background muscle activity. A one-way repeated-measures ANOVA revealed no significant difference among the ten conditions that included a perturbation (i.e., the perturbation-only and the nine perturbation-and-LAS conditions), either in elbow displacement at 50 ms (F(4.28, 47.03) = 1.152, p = 0.428, ηp² = 0.095) or in Vmax (F(9, 99) = 1.366, p = 0.428, ηp² = 0.110). Accordingly, no post hoc comparisons were performed. Furthermore, to assess whether LAS evoked an overt startle response during the experiment, the peak amplitude of the SCM EMG in the 20–120 ms window after LAS onset was compared with that in the pre-LAS baseline window. No statistically significant increase was detected (*t* (11) = 0.360, *p* = 0.726, *d*_z_ = 0.104). Thus, the SCM analysis provided no evidence of an overt startle response within the tested window, consistent with habituation across repeated LAS presentations.

Figure 3 illustrates the BB EMG waveforms of a representative participant across all nine LAS timings (A) and the corresponding group-averaged facilitation waveforms (B). To assess whether LAS produced supralinear facilitation of the LLR, each perturbation-and-LAS response was compared with the linear sum of the perturbation-only and LAS-only responses. In the representative participant, the perturbation-and-LAS response (blue) clearly exceeded the linear sum (black) within the LLR at LAS timings from 10 to 70 ms, whereas no such excess was evident within the SLR. In addition, at LAS timings of 50 ms and later, the facilitated portion of the response shifted progressively later in step with the LAS timing. A similar facilitation pattern was also observed in the group-averaged waveforms (perturbation-and-LAS minus linear sum).

**Fig 3.**
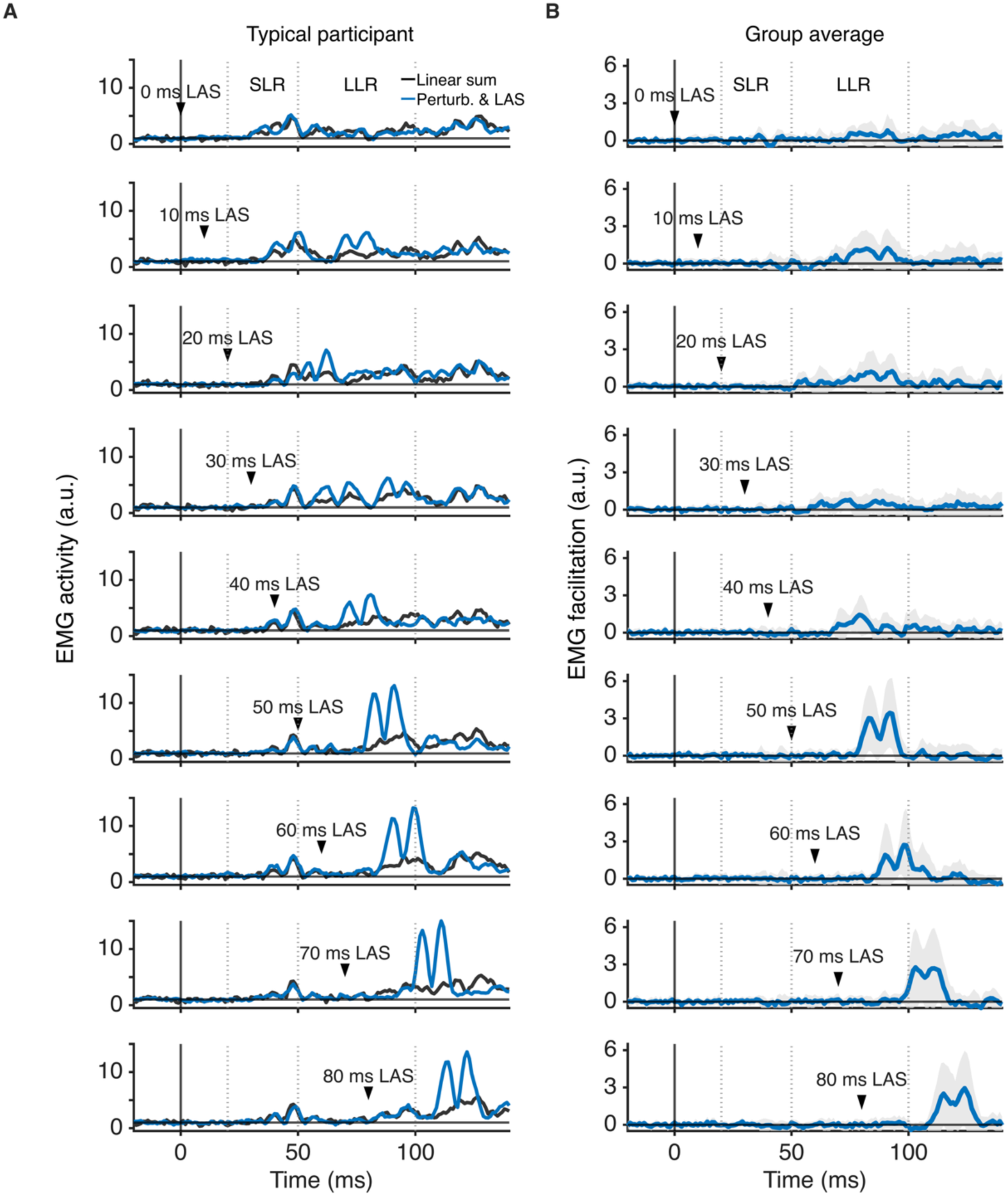
Supralinear facilitation of the stretch reflex in the biceps brachii (BB) muscle by LAS: EMG waveforms in a representative participant and the group average. **(A)** The BB EMG waveforms for a typical participant across all nine LAS timings. Blue line, perturbation-and-LAS condition; black line, linear sum (perturbation-only + time-shifted LAS-only − 1). Dotted vertical lines indicate boundaries of the SLR (20–50 ms) and LLR (50–100 ms) windows; arrowheads indicate LAS onset. EMG is normalised to the mean baseline (100-ms window before perturbation onset) and expressed in baseline-normalised units (a.u.), such that 1 corresponds to baseline activity. **(B)** Group-averaged BB EMG facilitation waveforms (perturbation-and-LAS minus linear sum) for each LAS timing; blue line, mean; grey shading, ± SD.

Figure 4A and 4B illustrate the group-mean facilitation and the corresponding one-sample *t*-value map (facilitation versus zero), respectively, over post-perturbation time and LAS timing, with the significant cluster outlined. To assess facilitation across the SLR and LLR windows, the facilitation map was submitted to a two-dimensional cluster-based permutation test. The test revealed a single significant cluster (*p* < 0.001, T-sum = 576.7), with no further cluster attaining significance. Significant facilitation was absent throughout the SLR window but present from the LLR window onward. At LAS timings of 0–50 ms, the facilitated interval remained within the predefined LLR window. From LAS timings of 50 ms and later, the facilitated window shifted together with LAS onset and progressively extended beyond the LLR, preserving an approximately stable interval relative to LAS onset (Fig. 4A, B).

**Fig 4.**
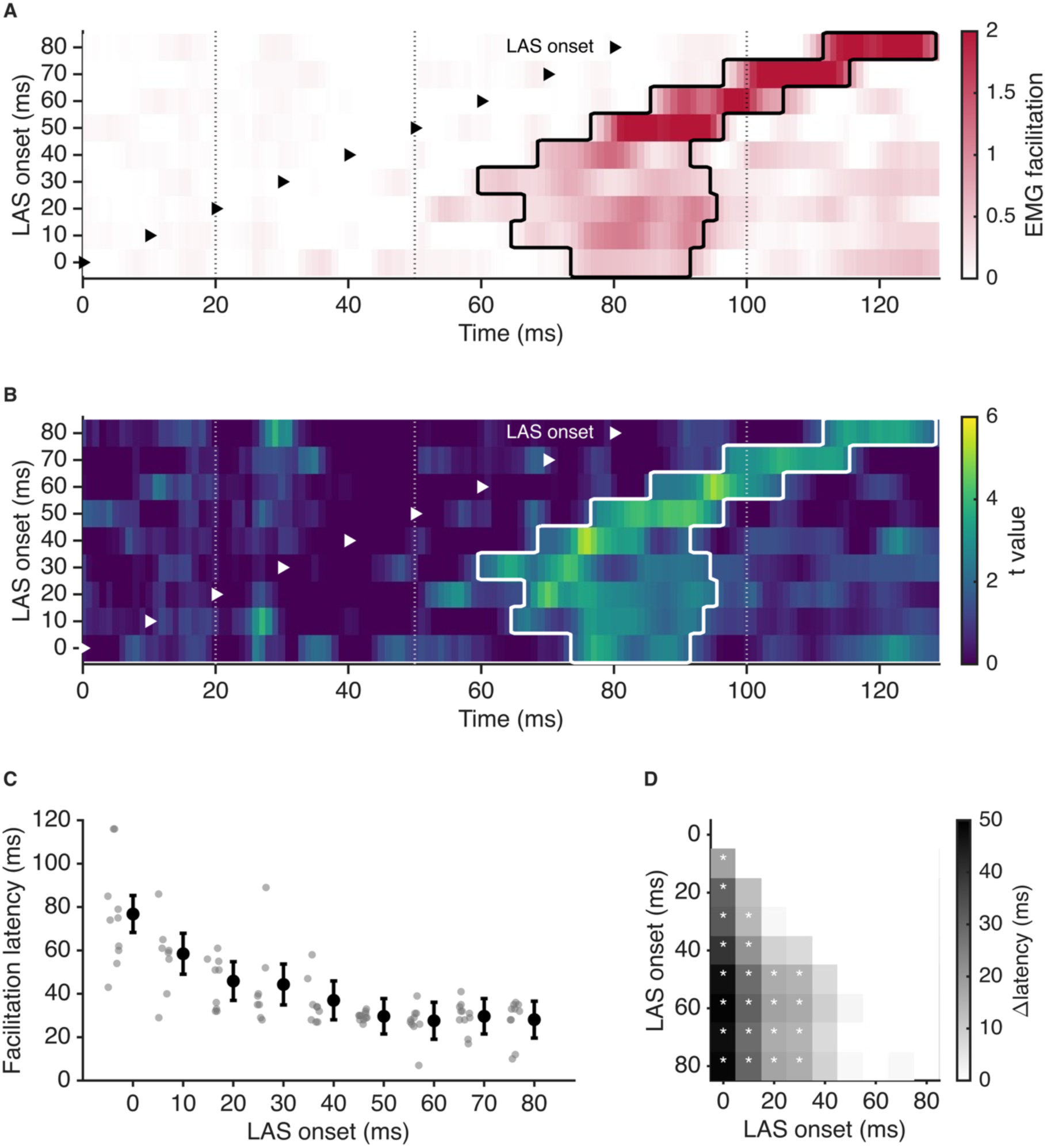
Cluster-based permutation test of LAS-induced facilitation and the facilitation latency across LAS timings. **(A)** Group-mean facilitation (observed − linear sum) shown as a heatmap (colour, facilitation magnitude in baseline-normalised units) over post-perturbation time (x-axis) and LAS timing (y-axis), after 5-ms smoothing along the time axis. The black contour outlines the significant cluster from the two-dimensional cluster-based permutation test. Dotted vertical lines mark 20, 50 and 100 ms (the SLR and LLR boundaries), and the diagonal markers indicate LAS onset for each row. **(B)** Corresponding one-sample *t*-value map (facilitation versus zero) with the same cluster contour; colour shows the group *t* value for the positive (facilitation) direction only, consistent with the one-sided test. **(C)** Facilitation latency (facilitation onset − LAS timing) as a function of LAS onset; grey dots, individual participants; black symbols, estimated marginal mean ± 95% CI from the linear mixed model. Because a facilitation onset was not detected in every participant at every LAS timing, the number of individual data points (grey dots) varies across LAS timings (see Table 1). **(D)** Pairwise differences in the estimated marginal latency between LAS timings (rows and columns, LAS onset). Greyscale encodes the signed difference in estimated marginal latency (earlier LAS timing minus later LAS timing; Δlatency, in ms), with values at or below zero shown in the lightest shade. Asterisks indicate contrasts that remained significant after Benjamini–Hochberg correction (*q* < 0.05); FDR-adjusted *q* values for all 36 contrasts are reported in Table S1.

**Table 1.** Facilitation latency per LAS timing.

| LAS onset (ms) | *n* | EMM [95% CI] (ms) | Median [IQR] (ms) |
| --- | --- | --- | --- |
| 0 | 10 | 76.8 [68.3, 85.3] | 74.5 [60.0, 85.0] |
| 10 | 8 | 58.5 [49.0, 67.9] | 59.5 [48.0, 63.0] |
| 20 | 9 | 45.9 [36.9, 54.8] | 51.0 [32.8, 55.3] |
| 30 | 8 | 44.3 [34.9, 53.7] | 37.0 [32.0, 46.0] |
| 40 | 9 | 37.0 [28.0, 45.9] | 34.0 [27.8, 38.0] |
| 50 | 11 | 29.6 [21.5, 37.8] | 30.0 [28.3, 30.8] |
| 60 | 10 | 27.6 [19.1, 36.1] | 27.5 [26.0, 31.0] |
| 70 | 11 | 29.6 [21.5, 37.8] | 31.0 [28.0, 33.5] |
| 80 | 10 | 28.1 [19.6, 36.6] | 32.5 [28.0, 35.0] |

Latency = facilitation onset (relative to perturbation onset) – LAS timing. EMM [95% CI], estimated marginal mean and 95% confidence interval from the linear mixed model; median [IQR], model-free per-timing median and interquartile range. *n*, number of participants contributing a detectable facilitation onset at that timing; a detectable onset did not occur in every participant at every LAS timing.

Figure 4C illustrates the relationship between LAS timing and the facilitation latency (facilitation onset minus LAS timing), including data from each participant (grey dots) together with the estimated marginal mean and its 95% confidence interval from the linear mixed model (black symbols). Figure 4D and Table 1 further summarise the pairwise differences in this latency among LAS timings. To test whether the facilitation latency differed among LAS timings, it was analysed with a linear mixed model. The facilitation latency depended significantly on LAS timing (*F* (8, 66.9) = 17.86, *p* < 0.001; marginal *R*^2^ = 0.585, conditional *R*^2^ = 0.656). Indeed, the facilitation latency was the longest at the LAS timing of 0 ms and shortened as LAS was delayed. At LAS timings of 50 ms and later, the EMMs ranged from 27.6 to 29.6 ms (Table 1; Fig. 4C). Pairwise contrasts (Benjamini– Hochberg corrected) showed that the facilitation latency at each of the LAS timings from 0 to 30 ms was significantly longer than those at each of the 50–80 ms timings, whereas no pairwise difference emerged among the 50–80 ms timings. The 40-ms timing was transitional: it differed significantly from the two earliest timings (0 and 10 ms) but not from either the adjacent 20–30 ms or the 50–80 ms timings (Fig. 4D). The model-free per-timing medians showed a qualitatively similar descriptive pattern across the 50–80 ms timings (27.5–32.5 ms; Table 1).

## Discussion

We combined mechanical perturbations with LAS across nine stimulus timings to test whether LAS-evoked reticulospinal drive facilitates the LLR. The principal findings were as follows: 1) LAS produced significant timing-dependent facilitation within the LLR, whereas no significant facilitation was detected in the SLR; 2) Such LLR facilitation shifted progressively later with a fixed delay of approximately 30 ms, when applying LAS at 50 ms or more after perturbation onset. On the basis of previously revealed data showing LAS-induced engagement of brainstem circuits including the reticular formation, we could regard the present data as evidence for an online contribution of the reticulospinal tract to the human stretch reflex, the fastest feedback mechanism for human motor control.

### LAS-induced facilitation specific to the LLR

A key feature of the present results is that LAS significantly facilitated the LLR, whereas no significant facilitation was detected in the SLR. This distinction is important because the SLR is primarily a spinal reflex component mediated by Ia afferents and spinal motoneurons. The facilitation pattern induced by LAS resembles that produced when TMS was superimposed on the stretch reflex, such that TMS did not facilitate the SLR but produced supralinear facilitation in the LLR (Day et al., 1991; Palmer and Ashby, 1992). This pattern supports the interpretation that similar to TMS, LAS provides a facilitatory effect confined to the reflex epochs to which supraspinal inputs are expected to contribute, rather than a general amplification of the stretch reflex.

The facilitated epoch shifted progressively later as LAS onset was delayed relative to perturbation onset. For LAS delivered at 50 ms or later, detectable facilitation emerged after an approximately constant latency from LAS onset (EMMs, 27.6–29.6 ms; Table 1; Fig. 4C, D). Thus, the facilitation was locked to LAS onset rather than occurring at a fixed time after perturbation onset. This fixed delay would serve as evidence that observed facilitation of LLR is attributable to the descending drive from the same origin, when applying LAS at 50 ms or more after perturbation onset. Although the latency pattern alone does not identify the pathway, and a contribution from the corticospinal tract or other descending pathways cannot be excluded, reticulospinal mediation is the most plausible interpretation because LAS strongly engages pontomedullary reticular circuits (Yeomans et al., 2002; Zheng and Schmid, 2023). Together with the selective facilitation of the LLR, this LAS-locked temporal shift supports an online reticulospinal contribution to the human LLR.

Previous studies have provided evidence for an offline contribution of reticulospinal drive to the LLR. Foysal et al. (2016) demonstrated that repeated pairing of brief loud auditory clicks (110 dB SPL) and peripheral muscle stimulation induced offline plasticity in LLR gain. Related paired-stimulation work has provided additional evidence for subcortical plasticity using StartReact and TMS-derived measures (Germann and Baker, 2021). The present study extends previous offline evidence by showing that LAS-evoked input provided online drive to the human LLR. Together with evidence from TMS studies (Day et al., 1991; Palmer and Ashby, 1992; Pruszynski et al., 2011; Perenboom et al., 2015), our findings support the view that the LLR is generated through coordinated contributions from multiple descending pathways, including corticospinal and reticulospinal pathways. The present study does not determine whether these pathways serve identical or distinct functional roles during the same LLR epoch. Rather, we believe that it provides a framework for testing that question in future work.

### Alternative explanations for LAS-induced LLR facilitation

We should consider other potential mechanisms behind the observed LAS-induced facilitation of LLR together with the reticulospinal interpretation. First, because LAS reliably activates startle-related brainstem circuits, the LLR facilitation could in principle be attributed to an acoustic startle response. However, this explanation is difficult to reconcile with the present results. In the LAS-only condition, SCM EMG activity in the 20–120 ms window after LAS onset did not differ from the pre-LAS baseline. Because SCM activity has been commonly used as a marker of overt startle responses (Brown et al., 1991; Valls-Solé et al., 1999), the present SCM analysis provided no evidence of an overt startle response within the tested window, supporting habituation across repeated LAS presentations. In addition, the present facilitation analysis compared the perturbation-and-LAS response with the linear sum of the perturbation-only and LAS-only responses. If LAS simply evoked a startle EMG response that was added to the perturbation-only EMG response, that activity would already be captured in the linear sum and would not appear as supralinear facilitation. Most importantly, the facilitation was too early to be startle-evoked muscle activity. For LAS delivered at 50 ms or later after perturbation onset, it began 27.6–29.6 ms after LAS, whereas the earliest component of the generalised startle reflex, in the sternocleidomastoid muscle, has a median latency of 58.3 ms (Brown et al., 1991; Carlsen et al., 2011). Thus, startle activity in an upper-limb muscle would be expected later still.

Second, because LAS is a startling stimulus and participants were prepared to correct the perturbation, a StartReact mechanism must also be considered. StartReact refers to the early release of a prepared motor response triggered by a startling stimulus. In upper-limb muscles, StartReact-related EMG activity has been reported to emerge approximately 72–108 ms after the LAS (Valls-Solé et al., 1999; Alibiglou and MacKinnon, 2012; Nonnekes et al., 2014; Castellote and Kofler, 2018; Walker et al., 2024). When LAS was delivered at perturbation onset (i.e., 0 ms LAS), the facilitation latency was 76.8 ms (95% CI, 68.3–85.3), which overlaps the range of StartReact-related activity. A StartReact contribution therefore cannot be excluded in that condition. For LAS delivered at 10 ms or later after perturbation onset, however, the facilitation latency was 58.5 ms or shorter. Moreover, for LAS delivered at 50 ms or later after perturbation onset, the facilitation latency remained constant at 27.6–29.6 ms. This was approximately half that shortest value. StartReact could therefore have contributed only when LAS was delivered at perturbation onset, and did not account for the main findings of the present study (i.e., LLR facilitation induced by LAS delivered at 50 ms or later after perturbation onset).

Third, the increase in arousal levels induced by LAS may have led to the present findings since the startle reflex is closely linked to arousal and emotional systems (Yeomans et al., 2002; Zheng and Schmid, 2023). However, three observations make a substantial contribution from these systems unlikely, as follows: 1) As the startle reflex is known to habituate within two to six presentations (Brown et al., 1991), the present participants were considered to be familiarised with LAS before the main experiment, which could indeed be supported by no SCM response during the experiment; 2) As noted above, for LAS delivered at 50 ms or later after perturbation onset, the LLR facilitation began 27.6–29.6 ms after LAS, which was earlier than the earliest component of the generalised startle reflex (sternocleidomastoid, median 58.3 ms; Brown et al., 1991); and 3) the SLR was not facilitated regardless of the timing of LAS application, whereas a general rise in arousal, motoneuron excitability, or reflex gain would be expected to facilitate the SLR as well. Although we do not exclude an effect of LAS on arousal or emotional systems, these systems do not appear to be critical for the LLR-selective facilitation observed here.

### Limitations and future directions

This study was limited to the biceps brachii muscle during elbow perturbation. Therefore, it remains unclear whether the present paradigm, which combines LAS and mechanical perturbation, would reveal comparable LLR facilitation when applied to other muscles or joints. A particularly important factor is that the strength and pattern of reticulospinal effects vary across muscles (Baker, 2011). Consequently, it cannot be assumed that the facilitation observed in BB will generalise to distal muscles, extensors or muscles acting at other joints. Additionally, the appropriate time lag between LAS and mechanical perturbation for selective facilitation of the LLR might differ across body parts. Although this limitation does not diminish the utility of the present method, if the stimulation technique developed in this study is to be extended to other body parts, the paradigm must be appropriately adjusted prior to experimentation.

Finally, one important question that remains open is whether the reticulospinal drive contributes to task-dependent modulation of the LLR. The LLR is strongly shaped by behavioural goals and task demands (Pruszynski et al., 2008), and the mechanisms underlying this flexibility may differ across task contexts. Indeed, cortical involvement has been demonstrated for modulation by environmental mechanics, whereas instruction-related modulation can persist during cortical suppression (Shemmell et al., 2009; Kurtzer, 2015). Future studies could combine the LAS paradigm developed in this study with different task conditions to test whether reticulospinal drive contributes to task-dependent LLR modulation, and whether its role overlaps with or differs from that of corticospinal drive. Therefore, the LAS paradigm developed in the present study, when used in conjunction with TMS, could serve as a useful framework for non-invasively examining the functional roles of descending pathways in rapid feedback control in humans.

## Conclusion

In summary, we used a supralinear summation paradigm combining mechanical perturbation and LAS to test whether LAS-evoked reticulospinal drive contributes online to the human LLR. LAS produced significant timing-dependent supralinear facilitation in the LLR, whereas no significant facilitation was detected in the SLR. This pattern indicates that LAS-evoked descending drive was integrated into the reflex response as it arrived within the LLR window. Together with prior evidence for corticospinal involvement in the LLR, the present findings support the view that rapid feedback control in humans is shaped by coordinated contributions from multiple descending pathways, including corticospinal and reticulospinal pathways. Beyond these specific findings, combining LAS with mechanical perturbation provides a non-invasive approach for probing reticulospinal function during human movement.

## Data Availability Statement

The data and MATLAB analysis code that support the findings of this study are openly available in Zenodo at https://doi.org/10.5281/zenodo.21867346.

## Author Contributions

H.S., D.N. and J.U. conceived and designed the study. H.S. performed the experiments and analysed the data. H.S., D.N. and J.U. interpreted the results. H.S. drafted the manuscript, and D.N. and J.U. revised it critically for important intellectual content. J.U. supervised the project. All authors have read and approved the final version of this manuscript and agree to be accountable for all aspects of the work in ensuring that questions related to the accuracy or integrity of any part of the work are appropriately investigated and resolved. All persons designated as authors qualify for authorship, and all those who qualify for authorship are listed. The experiments were performed at Saiseikai Higashikanagawa Rehabilitation Hospital, Japan.

## Generative AI Statement

During preparation of this manuscript, the authors used Anthropic Claude Code (versions 2.1.177–2.1.209; models claude-opus-4-8, claude-sonnet-5, claude-haiku-4-5-20251001, and claude-fable-5; accessed June–July 2026) and OpenAI Codex CLI (version 0.144.5; model gpt-5.6-sol; accessed July 2026) for English-language editing, assessment of manuscript structure and internal consistency, review of statistical reporting, and auditing of analysis code and project documentation. The authors critically reviewed and verified all suggestions and outputs. All scientific decisions were made by the authors, who take full responsibility for the final content of the manuscript.

## Acknowledgements

We thank Ms. Tomomi Hamaoka for her secretarial assistance and all members of our laboratory for their insightful comments on this work. We also thank Dr Hiroki Ebata, President of Saiseikai Higashikanagawa Rehabilitation Hospital, and the hospital staff for their support in conducting the experiments.

## Translational Perspective

The long-latency reflex (LLR) depends on descending motor pathways, yet the real-time contribution of the reticulospinal tract has remained inaccessible in humans. By combining mechanical perturbation with LAS, the present approach provides a non-invasive means of probing this contribution online, during movement. Because the reticulospinal tract is increasingly recognised as an important substrate for motor recovery after stroke and spinal cord injury, particularly when corticospinal function is compromised, a non-invasive and movement-compatible assay of its function could become a valuable tool for future research. It may support the assessment and tracking of descending pathway function in neurological patients and help guide rehabilitation strategies that engage reticulospinal circuits.

## Supporting Information

**Table S1.**
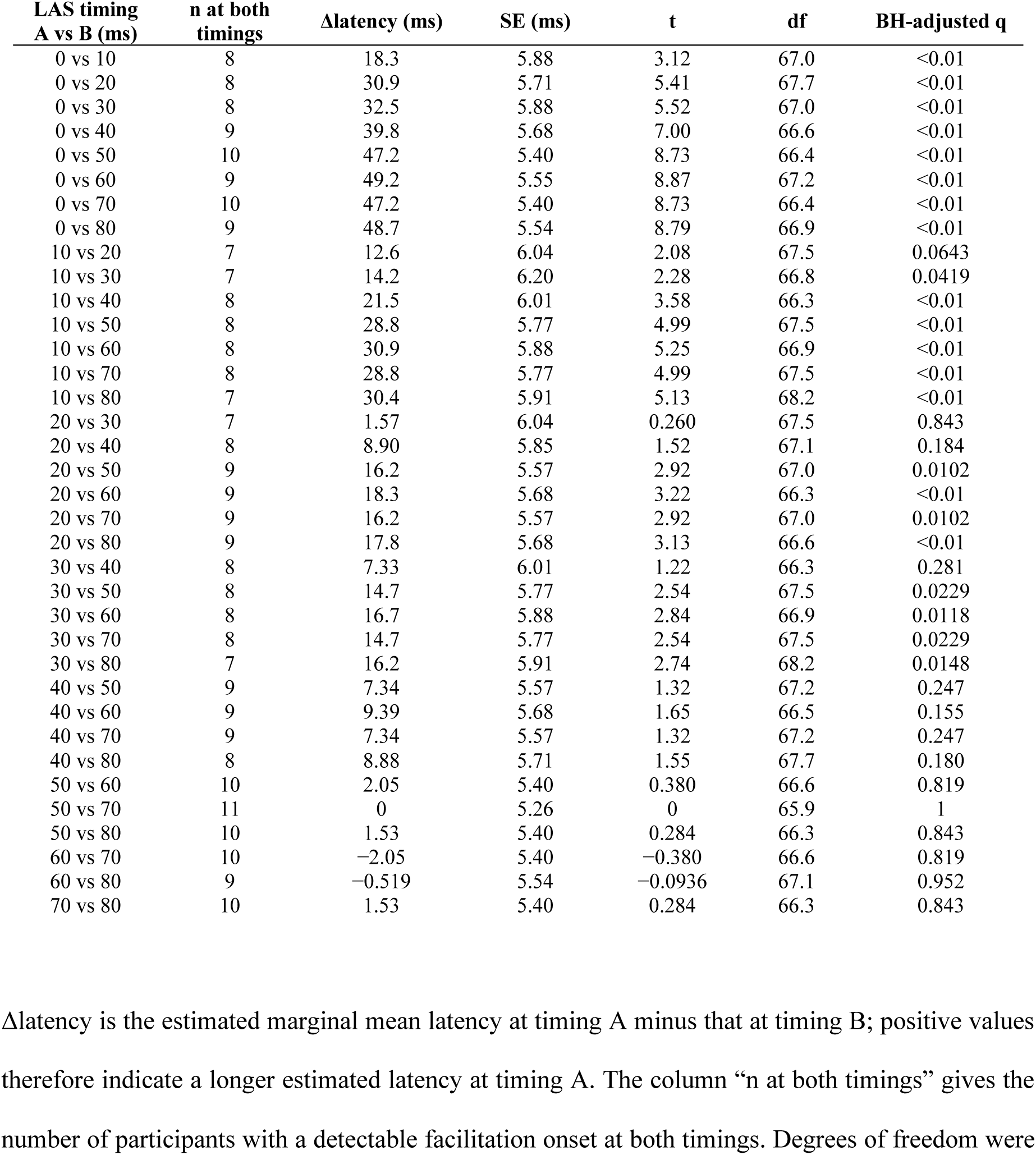
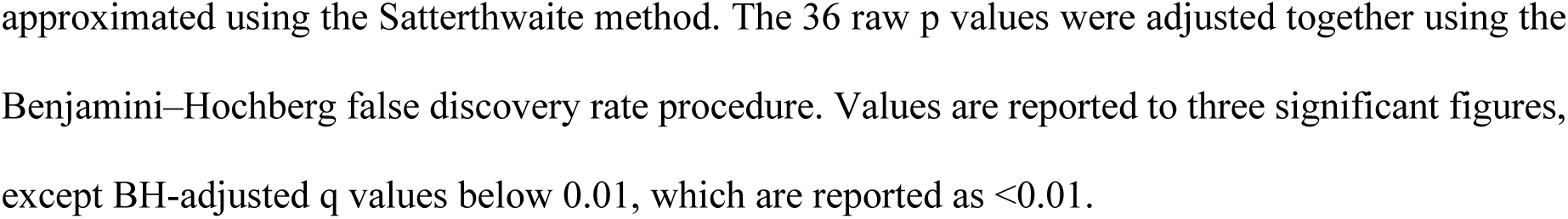
Pairwise contrasts of estimated marginal mean facilitation latency between LAS timings.

| LAS timing<br>A vs B (ms) | n at both<br>timings | Δlatency (ms) | SE (ms) | t | df | BH-adjusted q |
| --- | --- | --- | --- | --- | --- | --- |
| 0 vs 10 | 8 | 18.3 | 5.88 | 3.12 | 67.0 | <0.01 |
| 0 vs 20 | 8 | 30.9 | 5.71 | 5.41 | 67.7 | <0.01 |
| 0 vs 30 | 8 | 32.5 | 5.88 | 5.52 | 67.0 | <0.01 |
| 0 vs 40 | 9 | 39.8 | 5.68 | 7.00 | 66.6 | <0.01 |
| 0 vs 50 | 10 | 47.2 | 5.40 | 8.73 | 66.4 | <0.01 |
| 0 vs 60 | 9 | 49.2 | 5.55 | 8.87 | 67.2 | <0.01 |
| 0 vs 70 | 10 | 47.2 | 5.40 | 8.73 | 66.4 | <0.01 |
| 0 vs 80 | 9 | 48.7 | 5.54 | 8.79 | 66.9 | <0.01 |
| 10 vs 20 | 7 | 12.6 | 6.04 | 2.08 | 67.5 | 0.0643 |
| 10 vs 30 | 7 | 14.2 | 6.20 | 2.28 | 66.8 | 0.0419 |
| 10 vs 40 | 8 | 21.5 | 6.01 | 3.58 | 66.3 | <0.01 |
| 10 vs 50 | 8 | 28.8 | 5.77 | 4.99 | 67.5 | <0.01 |
| 10 vs 60 | 8 | 30.9 | 5.88 | 5.25 | 66.9 | <0.01 |
| 10 vs 70 | 8 | 28.8 | 5.77 | 4.99 | 67.5 | <0.01 |
| 10 vs 80 | 7 | 30.4 | 5.91 | 5.13 | 68.2 | <0.01 |
| 20 vs 30 | 7 | 1.57 | 6.04 | 0.260 | 67.5 | 0.843 |
| 20 vs 40 | 8 | 8.90 | 5.85 | 1.52 | 67.1 | 0.184 |
| 20 vs 50 | 9 | 16.2 | 5.57 | 2.92 | 67.0 | 0.0102 |
| 20 vs 60 | 9 | 18.3 | 5.68 | 3.22 | 66.3 | <0.01 |
| 20 vs 70 | 9 | 16.2 | 5.57 | 2.92 | 67.0 | 0.0102 |
| 20 vs 80 | 9 | 17.8 | 5.68 | 3.13 | 66.6 | <0.01 |
| 30 vs 40 | 8 | 7.33 | 6.01 | 1.22 | 66.3 | 0.281 |
| 30 vs 50 | 8 | 14.7 | 5.77 | 2.54 | 67.5 | 0.0229 |
| 30 vs 60 | 8 | 16.7 | 5.88 | 2.84 | 66.9 | 0.0118 |
| 30 vs 70 | 8 | 14.7 | 5.77 | 2.54 | 67.5 | 0.0229 |
| 30 vs 80 | 7 | 16.2 | 5.91 | 2.74 | 68.2 | 0.0148 |
| 40 vs 50 | 9 | 7.34 | 5.57 | 1.32 | 67.2 | 0.247 |
| 40 vs 60 | 9 | 9.39 | 5.68 | 1.65 | 66.5 | 0.155 |
| 40 vs 70 | 9 | 7.34 | 5.57 | 1.32 | 67.2 | 0.247 |
| 40 vs 80 | 8 | 8.88 | 5.71 | 1.55 | 67.7 | 0.180 |
| 50 vs 60 | 10 | 2.05 | 5.40 | 0.380 | 66.6 | 0.819 |
| 50 vs 70 | 11 | 0 | 5.26 | 0 | 65.9 | 1 |
| 50 vs 80 | 10 | 1.53 | 5.40 | 0.284 | 66.3 | 0.843 |
| 60 vs 70 | 10 | -2.05 | 5.40 | -0.380 | 66.6 | 0.819 |
| 60 vs 80 | 9 | -0.519 | 5.54 | -0.0936 | 67.1 | 0.952 |
| 70 vs 80 | 10 | 1.53 | 5.40 | 0.284 | 66.3 | 0.843 |

Δlatency is the estimated marginal mean latency at timing A minus that at timing B; positive values therefore indicate a longer estimated latency at timing A. The column “n at both timings” gives the number of participants with a detectable facilitation onset at both timings. Degrees of freedom were
approximated using the Satterthwaite method. The 36 raw p values were adjusted together using the Benjamini–Hochberg false discovery rate procedure. Values are reported to three significant figures, except BH-adjusted q values below 0.01, which are reported as $<0.01$ .

